# HistoFlow: Label-Efficient and Interactive Deep Learning Cell Analysis

**DOI:** 10.1101/2020.07.15.204891

**Authors:** Tim Henning, Benjamin Bergner, Christoph Lippert

**Author notes:** Correspondence: *Tim Henning, Benjamin Bergner or Christoph Lippert.

## Abstract

Instance segmentation is a common task in quantitative cell analysis. While there are many approaches doing this using machine learning, typically, the training process requires a large amount of manually annotated data. We present HistoFlow, a software for annotation-efficient training of deep learning models for cell segmentation and analysis with an interactive user interface.

It provides an assisted annotation tool to quickly draw and correct cell boundaries and use biomarkers as weak annotations. It also enables the user to create artificial training data to lower the labeling effort. We employ a universal U-Net neural network architecture that allows accurate instance segmentation and the classification of phenotypes in only a single pass of the network. Transfer learning is available through the user interface to adapt trained models to new tissue types.

We demonstrate HistoFlow for fluorescence breast cancer images. The models trained using only artificial data perform comparably to those trained with time-consuming manual annotations. They outperform traditional cell segmentation algorithms and match state-of-the-art machine learning approaches. A user test shows that cells can be annotated six times faster than without the assistance of our annotation tool. Extending a segmentation model for classification of epithelial cells can be done using only 50 to 1500 annotations.

Our results show that, unlike previous assumptions, it is possible to interactively train a deep learning model in a matter of minutes without many manual annotations.

## Introduction

Nucleus segmentation is an important topic in quantitative cell analysis and forms the foundation for a variety of analysis tasks, e.g., automated cell movement tracking (1). It is a well/studied topic and much research has been done to achieve good segmentation results for different imaging techniques. In our case, we will evaluate the presented tool on fluorescence breast cancer images.

Recent studies showed that deep learning could lead to more accurate results than traditional algorithms and shallow learning techniques such as Random Forests (2) (3).

While many tools allow the application of neural networks to microscopy images and some even provide a simple to use model repository for it (4) (5) (6) (7), to our knowledge, there is currently no tool that enables users to train a deep neural network in the same integrated environment.

This could be due to the fact that deep learning requires large amounts of annotated training data. Creating them by letting experts annotate microscopy images is time/consuming and expensive. For example, it took 50 hours to annotate the data in one of the above/mentioned studies (2). In some cases, it may not even be possible to create such a dataset because of time or data restrictions. Relying on already available annotated datasets would limit research to their specific modalities.

While interactivity in the form of quick results, fast iteration cycles and an integrated workflow can simplify research a lot, only shallow learning approaches such as in the *ilastik* (7) software offer it yet. There is also the potential to create a more intuitive annotation tool, as existing solutions such as *Caliban* from (6) have a steep learning curve.

### A. Key Contributions

We demonstrate that HistoFlow allows to

- annotate cells six times faster than without the assistance of its integrated annotation tool
- use synthetic training data to train deep neural network models in the matter of minutes, which perform comparably to those trained on real images
- train a per-cell classifier to detect epithelial cells using only 50 to 1500 annotations, partially from biomarkers as weak annotations
- use transfer learning to adapt a model trained with artificial data with additional manual annotations to work better for real fluorescence breast cancer images.

The node-based graphical user interface (GUI) allows interactive training and analysis, while the remote back end with accelerated deep learning using a universal U-Net (8) architecture and simplified Deep Watershed Transform (9) makes it easy to share results with other researchers.

We hereby address the issues of deep learning cell analysis pointed out in (2), like the need for manual annotations, longer preparation- and runtime, higher computational requirements and less transferability between datasets due to specific training data artifacts.

The source code of the software, including tutorials for model training and application, as well as pre-trained models are available on GitHub^1^ under the MIT license.

## Related Work

### B. Similar Tools

There are already many applications that try to solve the problem of cell segmentation using machine learning. This section describes the most popular ones and explains how our software differs from them.

#### B.1. Cell Profiler

According to citations, CellProfiler (10) is one of the most popular open-source tools for cell analysis tasks. It allows to create powerful image pipelines with traditional algorithms and shallow learning, e.g., decision trees (11). Since version 3.0, it also supports to apply Convolutional Neural Networks (CNNs). In contrast to this work, it does not allow to train them (4). Computational heavy tasks can be executed remotely, however, only using a commandline interface and not through the GUI as in this work.

#### B.2. ilastik

The focus of ilastik (12) is on interactive shallow learning, often with decision trees. It allows to apply deep learning models (7), too, and even supports an open model zoo, although it is not possible to train them in the application or even refine them interactively.

#### B.3. ImageJ Weka Plugin

The ‘Trainable Weka Segmentation’ plugin (13) for ImageJ (14) and its distribution Fiji allows to train and apply machine learning, but focuses on shallow learning. The newer DeepImageJ plugin (5) allows to apply deep learning models and features ready-to-use models for a variety of tasks such as segmentation, super-resolution and virtual staining. However, it does not support to train them or execute them on a remote GPU server.

#### B.4. DeepCell 2.0

Other than the previously mentioned tools, DeepCell 2.0 (15) is mainly used through a web interface. While it allows to train and apply deep learning models in a distributed cloud, the user interface is very limited at the moment. Compared to our tool, it also does not offer an integrated analysis environment.

### C. Segmentation Algorithms

This work uses the popular U-Net architecture (8) for semantic segmentation in combination with a simplified Deep Watershed Transform (DWT)(9) for instance localization. A similar approach was also used for the first place solution of the 2018 Data Science Bowl (16).

HoVer-Net (17) also combines segmentation and classification. Instead of DWT, it predicts horizontal and vertical gradients in the cells to find the boundaries between them. This idea is also found in the second place solution in (16).

Others use the proposal-based Mask R-CNN architecture (18). In contrast to our solution, it can’t distinguish two cells sharing the same bounding-box. Another promising - but not yet well studied - option is to extend U-Net with a recurrent neural network (19).

### D. Artificial Training Data Generation

Lehmussola et al.(20) created a MatLab-based framework to simulate fluorescence microscopy images to facilitate the validation of quantitative image processing techniques in 2007, however, it is not possible to create ground truth images suitable for the Deep Watershed Transform we are using.

Torrubia et al. (21) experimented with a Generative Adversarial Network (GAN) to generate synthetic training data in the 2018 Data Science Bowl. They transferred the style of a tissue type or imaging technique that they did not have enough ground truth for to images with a different style but with available ground truth. In a more recent paper (22), the pix2pixHD GAN was used to convert generated segmentation masks to synthetic images of red blood cells. While their results look promising and a GAN may be a good addition in the future, we propose a simpler solution that does not require any training to generate the images.

## Training Workflow

We present our training workflow for a model for simultaneous nucleus instance segmentation and classification into epithelial and other cells.

Traditionally, a large scale dataset with annotations of the individual nuclei and their phenotype would be required for this. To create it, one would have to segment the nuclei by hand and then mark each epithelial cell, which is a very time consuming and difficult task.

We propose a way to train such a model with only few manual annotations. It is divided into the following three phases that are shown in Fig. 1:

**Fig. 1.**
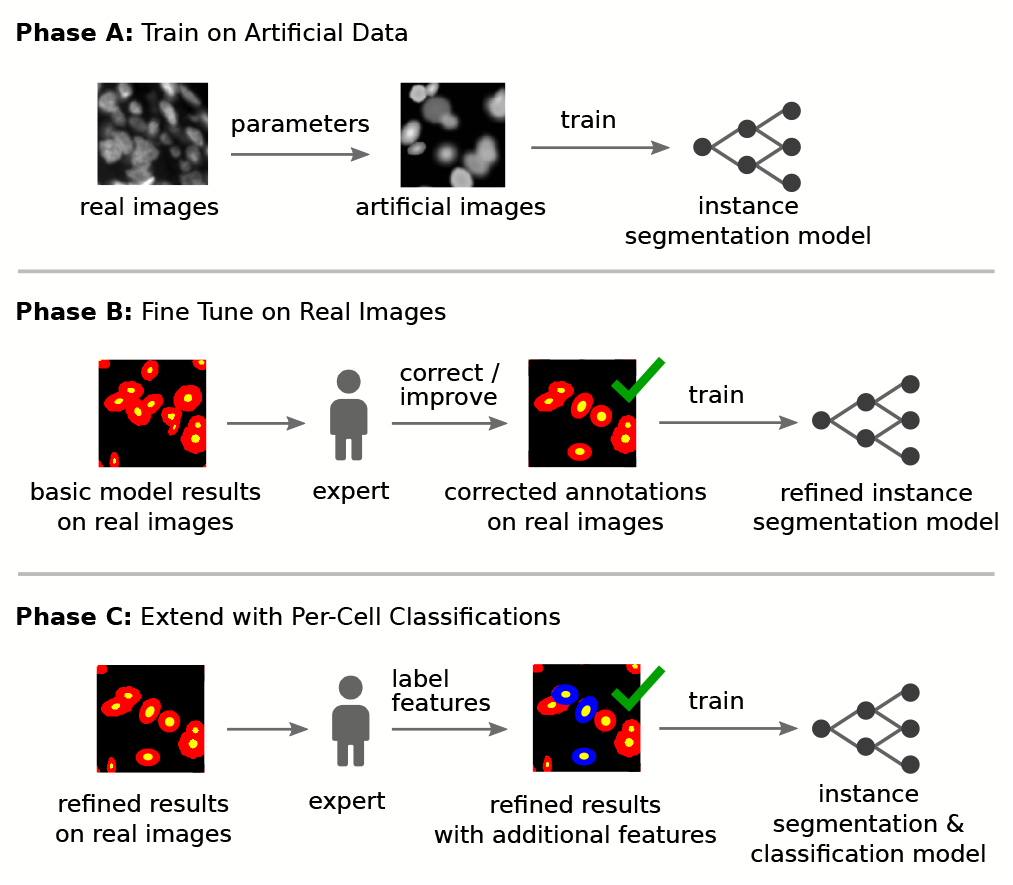
Training Workflow Example (real image patch in 1. row taken from *Image A*, patches in 2. and 3. row show the output of the neural network which is then processed using DWT)

### Phase A

Generate artificial training data - including their pixel-perfect generated ground truth results - with the help of parameters derived from the real images (see Section D). The neural network is then first trained using only the artificial data. Because the artificial data does not distinguish between any particular phenotype, this results in a model that is only able to do instance segmentation.

### Phase B (optional)

In the case the artificial training data does not exactly match the real images that have to be segmented, a second refinement step can be done. The first model will be applied to real images and errors in the resulting segmentation such as nuclei merges, splits and missed nuclei are corrected by an expert (see Section D). The corrected result is then used to further train the instance segmentation model. This transfer learning with a human-in-the-loop allows the usage of the artificial training data, which is at the moment limited to breast tissue cells on fluorescence images, for a wide range of similar looking tissue.

It is also possible to train a model with only manual annotations, in which case Phase A will be skipped and Phase B is mandatory.

### Phase C

To extend the model to also do per-cell tissue type classification at the same time, the model from phase A or B is applied to real images and the tissue type of a small number of cells is annotated by an expert (see section M for the actual number of cells). To simplify this step, weak annotations can be used, too. In this case, the intensity per cell of the Cytokeratin-19 staining is shown on a plot and using a single click all cells within a value range can be labeled as epithelial. After optionally hand/tuning those weak annotations, the annotations are used to train an additional channel of the previous model. The resulting model is then able to do instance segmentation and instance tissue type classification at the same time.

## Annotation Workflow

Our integrated annotation tool supports multiple modes of operation with varying levels of support that complement each other for different scenarios. In the above/presented example workflow, they are used for *Phase B* and C. Everything described for cells can also be applied to only the nuclei.

### Fully manual

A cell instance is created by clicking on the center of it. The outline is drawn by clicking there again and dragging the mouse on the edge of the cell, see fig. 2a. This can also be used to improve the outline of existing incorrect annotations. Cells can be deleted by a single click in this mode.

**Fig. 2.**
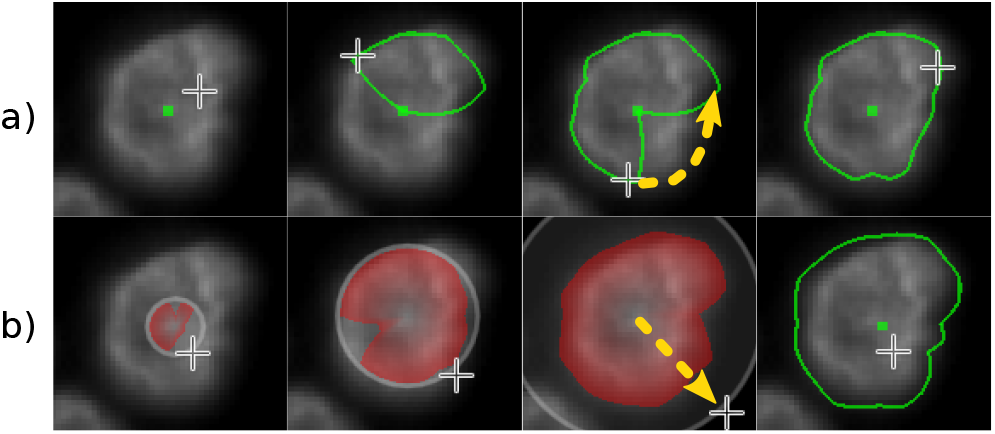
Annotation Modes: a) Fully Manual and b) Interactive Watershed. Yellow arrows illustrate the mouse movement. The Region Grow mode is not shown as it only requires marking the center of the nucleus as user input.

### Interactive Watershed

The user clicks on the center of a cell and opens a circular area around it by dragging the mouse. Within this range, an attempt is made to estimate the outline of the cell with a Watershed (23) inspired algorithm. Unlike the original algorithm, it does not flood the area starting from the local minima but finds the minimum value on a set of radial lines from the center to the edge of the area. The area of the cell is spanned between those points. The user sees the currently estimated outline while moving the mouse, hence the word ‘Interactive’ in the name, see fig. 2b.

To use this mode, an image channel must first be selected that provides the energy levels for the Watershed algorithm.

After two or more cells have been annotated, a simple click on the center of a cell is sufficient if the boundary is clearly visible. In this case, the software iterates over the range from the smallest to the largest existing cell radius, applies the Watershed algorithm described above to all of them and chooses the best fitting outline automatically.

### Region Grow

If there is already a semantic - but not instance - segmentation available, e.g., from a neural network model as in *Phase A* from the example workflow, it will be sufficient to first mark only the cell centers with a single click. In a second step, the outlines of the cells are found by a marker-based region grow algorithm (24). It either draws a region from the cell centers to the edge of the mask or until the region meets another region. The semantic segmentation makes it therefore faster to annotate the boundaries and is a simple form of machine learning assisted annotation.

### Labels from weak annotations

It is possible to plot the intensity of a staining per cell and mark all cells above a certain threshold as belonging to a specific category. By clicking on specific instances and overwriting the label, outliers can be fixed manually. This mode is useful to train a classifier based on a staining which is a good predictor for a certain biomarker, but that may not always be available due to time or cost restrictions.

## Artificial Data Creation

We present a way to generate artificial training data that simulates the DNA highlighting channel of a fluorescence image. It is a lightweight model-based technique that does not need any machine learning. The results can be seen in Fig. 3.

**Fig. 3.**
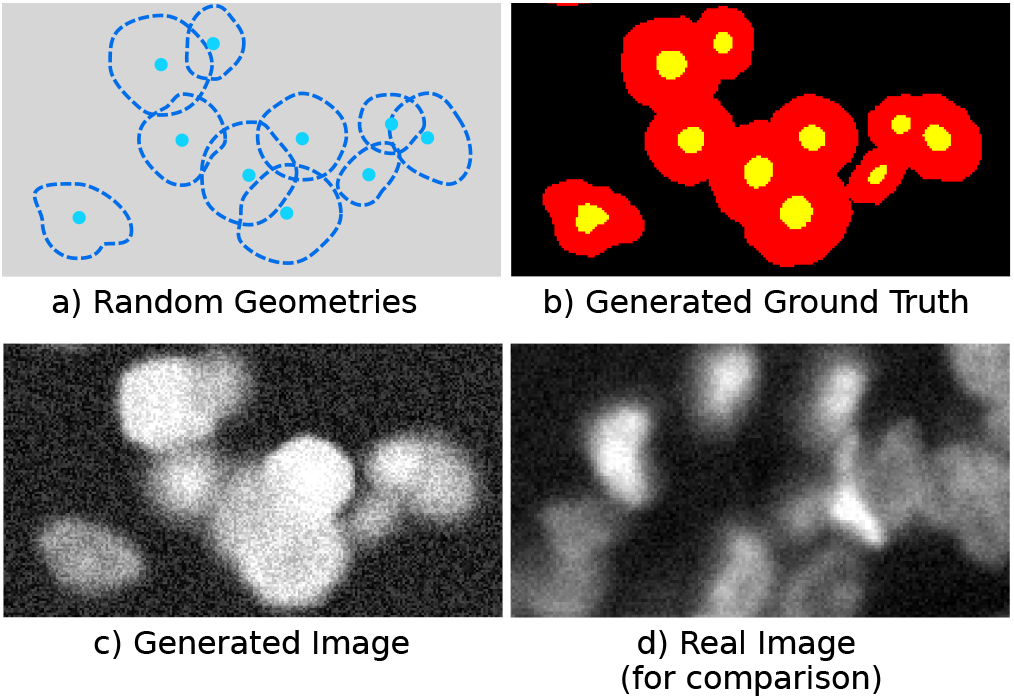
Different steps of artificial data generation. (a) is a visualization of the used geometries, (b) and (c) show the images generated from them and by comparing with (d), it can be seen how similar the result is to a real DAPI-stained breast tissue fluorescence image.

At first, the pixel resolution of the final image and the density of cells is specified by the user. The center coordinates of the nuclei are then randomly distributed on the image.

The shape of a nucleus is described as the offset of 24 equally distributed radii from the maximum radius, similar to (25). The maximum radius is assigned using a uniform random distribution between a minimum and maximum value. The range can be set numerically or the user annotates the smallest and largest expected nucleus on a real image.

The radii offsets are relative, normalized values that are uniformly random assigned between 0.9 and 1.0. With a probability provided by the user, the radii offsets of some nuclei are multiplied with an ellipsoid factor between 0.14 and 0.7 to generate elongated nuclei.

In addition, for each cell, the number of ‘twins’ is set by a uniformly random distribution between zero and an integral maximum twin count provided by the user. The number determines how many times the nucleus will be duplicated next to each other. This simulates cells right after they divided and, according to earlier experiments, helps to detect them correctly.

The nuclei are then drawn to an image by filling OpenGL meshes with random circular intensity gradients. At the moment, the range of gray levels is hard/coded to values typical for breast tissue fluorescence images that can be found in the provided code. A key insight was to invert some of the gradients to also detect nuclei, which appear darker in the middle due to being at the top of the tissue slice. Random, up to eight pixel large dots simulate dust and large, up to 400 pixel large objects with soft gradients simulate uneven lighting. The cells are blurred using a Gaussian blur GLSL shader with five pixel radius to simulate an offset to the focal plane. General image noise is added later during patch-wise preprocessing. A comparison between an artificial image and a real image can be seen in Fig. 3c.

The ground truth images are generated by first drawing the full cell area on the mask channel using OpenGL. The thinned nuclei for the second energy level of DWT reuse the same OpenGL objects but with a smaller scale factor and are drawn on the second image channel. An example ground truth patch can be seen in Fig. 3b.

## Node-Based User Interface and Client-Server Architecture

The front end is written platform/independently in C++ and QML using the Qt 5 framework. Representing algorithms, neural network models and images as connectable “blocks” in the software allows to compare them interactively. Evaluation metrics of different segmentations can be calculated in real/time to be able to tune hyperparameters and see how they affect the performance instantly. Missed (false negative, FN) and extra cells (false positive, FP) compared to the ground truth can be highlighted to understand the shortcomings of a model.

The back end is used to offload training and inference from the client. Images can be uploaded to the server to easily share experiment results with other researchers. The back end is written in Python using the Flask framework for REST communication and the fast.ai library (26) for GPU accelerated deep learning. The hardware used for the training and inference times reported in Section K consisted of an AMD Ryzen 3600 CPU, 8 GB system memory and one Nvidia RTX 2070 GPU with 8 GB graphics memory.

## Deep Learning Pipeline

### E. Neural Network Architecture

The used neural network architecture is the fully convolutional U-Net variant (27) from the fast.ai library, which is an improved version of the original one from Ronneberger et al. ResNet18 was chosen as the encoder.

For instance separation, the software uses Deep Watershed Transform (9). It is simplified so that the first channel of the output is the semantic segmentation mask and the second are the thinned nuclei that serve as the region seeds. It only uses those two channels as “energy” levels. Similar to (28), we do not predict unit vectors as they do not seem to improve the result a lot.

### F. Data

For development and validation, two fluorescence microscopy images from a breast cancer study were used (referred to as **Image A** and **Image B**). They measure about 9400 x 7600 pixels and show approximately 10’000 cells per image. Only the DNA-marking DAPI channel was used in the segmentation experiments. The Cytokeratin-19 (CK19) and Vimentin (VIM) stainings were used as weak annotations for the epithelial classification experiments.

To train networks without manual annotations, the **Artificial Dataset A** was generated. It contained a total of 40’000 cells on a single 14’000×14’000 pixel image (density was set to 10%). The noise augmentation parameter was set to 40% and the brightness change parameter to 75%. The minimum and maximum nucleus sizes were set through five example cells from *Image A*. The twin parameter was set to two.

In general, any images showing approximately round cells can be used with the software.

### G. Ground Truth

The ground truth for the real images was created by applying the best working neural network *Model D* (see L.3) to parts of *Image A* and *Image B* containing each about 3000 cells and correcting the segmentations by hand. As this may introduce a bias towards the specific way the neural network segments the cells, attention was paid to remove all of the characteristic errors of the model and to make the ground truth data as neutral and accurate as possible. The errors where mostly nuclei splits and missed nuclei, the outline was most of the time correct.

### H. Pre-Processing

The pre-processing is mostly done in the GUI. The user can adjust the black and white point and gamma curve of each image channel separately. These parameters are used for visualization as well as processing.

To get a set of training data, patches are cropped from random positions and with random 0-360° rotations equivalently from the input and target images. The patches from the input images will be randomly augmented with Gaussian noise and brightened or darkened by up to 50%.

### I. Post-Processing

Most of the post-processing can be done interactively in the GUI and depends on the actual use case. There are two operations that always have to be done in order to get the positions and boundaries of the nuclei. They are a simplified version of the Deep Watershed Transform with only two energy levels and no unit vectors:

First, finding the nuclei centers on the channel that is trained to output thinned nuclei. For this, a connected-components algorithm is used. Second, a region growing algorithm is applied to the semantic segmentation output channel with the nuclei centers as seeds.

Optionally, if classification is done, too, the results of it will be extracted from the output by averaging the pixels of each cell in the relevant output channel.

## Experiments

### J. Evaluation Metrics

For each ground truth (GT) nucleus, the nearest predicted nucleus will be marked as a true positive (TP) if the distance between their centers is smaller or equal the radius of the GT nucleus. Only one predicted nucleus can be marked as a true positive for each GT nucleus. The rest of the predicted nuclei are regarded as false positives (FP). The GT nuclei that have no predicted counterpart form the group of the false negatives (FN). The category of the true negatives (TN) is not applicable to the problem of finding one type of instances on an image; it is therefore always regarded as zero. Usually true positives for instance segmentation are defined as having an IoU greater than a certain threshold (see (2)). We differ from this approach to be able to calculate the metrics in real/time. This allows to tune post-processing parameters while observing how the evaluation metrics react. The data structure used to store the nuclei is similar to (25).

### K. Evaluation of Fast Annotations

One key aspect of the software is to help annotate cells faster and easier. In order to evaluate this, we asked three users to annotate the same 50 cells repeatedly with the different levels of support through our software and compared the time it takes to do this.

Beside that, we also asked them to improve an already existent segmentation result, so that the subjective quality becomes the same as with the fully manual annotations. This should simulate the real/world scenario where an instance-segmentation model is already available and one would only have to correct and fine/tune its results. This can be done either using the ‘fully manual’ or the ‘interactive Watershed’ mode. *Model C* from see Section L.3 was used for the initial segmentation.

Note that a forward pass of the network, including the upload of the image data to the back end, takes about 60 seconds for an image with 10’000 cells. Analyzing only the area of the 50 cells of this experiment takes about two seconds.

In the last step, the users were asked to correct the results of a traditional segmentation algorithm to see whether it really takes longer to correct those than the ones of the neural network. As an example, the result of the QiTissue segmentation result from section L.4 was used.

Of the three participants, two of them had no prior experience in the tool except for a short introduction and one was an experienced user. More support always led to less time required for the annotations. The largest difference between the participants was seen in the fully manual mode, where the experienced user was twice as fast as the others. All results can be seen in fig. 4.

**Fig. 4.**
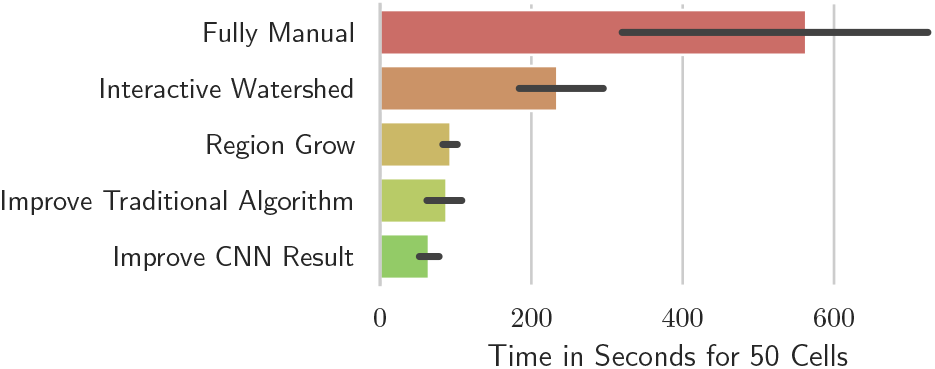
Time to annotate the boundary of 50 cells in seconds using different annotation modes.

### L. Evaluation of Segmentation Results

#### L.1. Artificial Data vs. Manual Annotations

To validate the quality of the synthetic data, two models were trained in the same way but one with only artificial data (**Model A**) and the other with the same amount of manual annotations (**Model B**). Both training data sets contained about 3600 cells. The manually annotated training data came from *Image* A. Both models were tested against a manually annotated part of *Image B* containing 3000 cells. The results are shown in fig. 6.

While the F1 scores did not show a large difference (Model A: 0.905, Model B: 0.914), the models had different strengths. The precision of the model with manual annotations was a lot better (0.957 vs. 0.886), but the recall was better for the model trained with artificial data (0.875 vs. 0.926).

#### L.2. Amount of Artificial Data

Although an almost unlimited amount of artificial data can be generated (the current implementation is limited to 40’000 cells), the following experiment shows how much training data is actually required. This is helpful in order to keep the training time short and allow fast iterations.

While keeping all other parameters the same, a neural network was trained using different subsets of the *Artificial Dataset A*. Each time an area containing the specified amount of cells was chosen and patches were randomly cut from this area. Each patch contained up to 30 cells and was randomly augmented with noise and brightness changes. The network was initialized with the same ImageNet weights in every case.

Figure 5 shows that between 10 and 500 unique cells, a significant increase in the resulting performance can be seen. Using more than 500 unique cells does not result in a large performance benefit. Only the use of 40’000 cells compared to 10’000 leads to an actual improvement again.

**Fig. 5.**
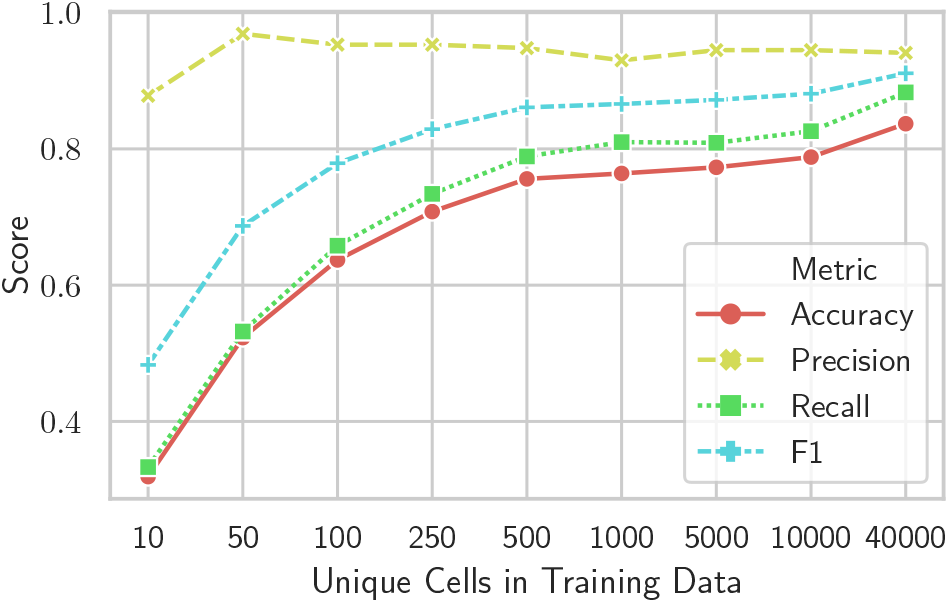
Segmentation quality with different amount of artificial training data. Evaluated on a test set containing about 3000 real cells from *Image B*.

Training with 500 cells and 800 patches will take about two minutes while training with 40’000 cells and 8000 patches will take about 20 minutes. These short training times highlight that deep learning can be integrated interactively in an analysis workflow.

#### L.3. Combination of Artificial Data and Manual Annotations

To test whether it is possible to further tune a model trained only with artificial data with additional manual annotations, the best model from the last section (called **Model C**) was further trained with the 3600 manually annotated cells used for *Model A*. This hybrid **Model D** was then tested using the manual annotations on *Image B*.

The results can be seen in fig. 6. *Model D* outperforms the models trained with only one kind of data. Its F1 score is the best that could be achieved on *Image B*, although the recall is lower than the one of the model trained only with artificial data.

**Fig. 6.**
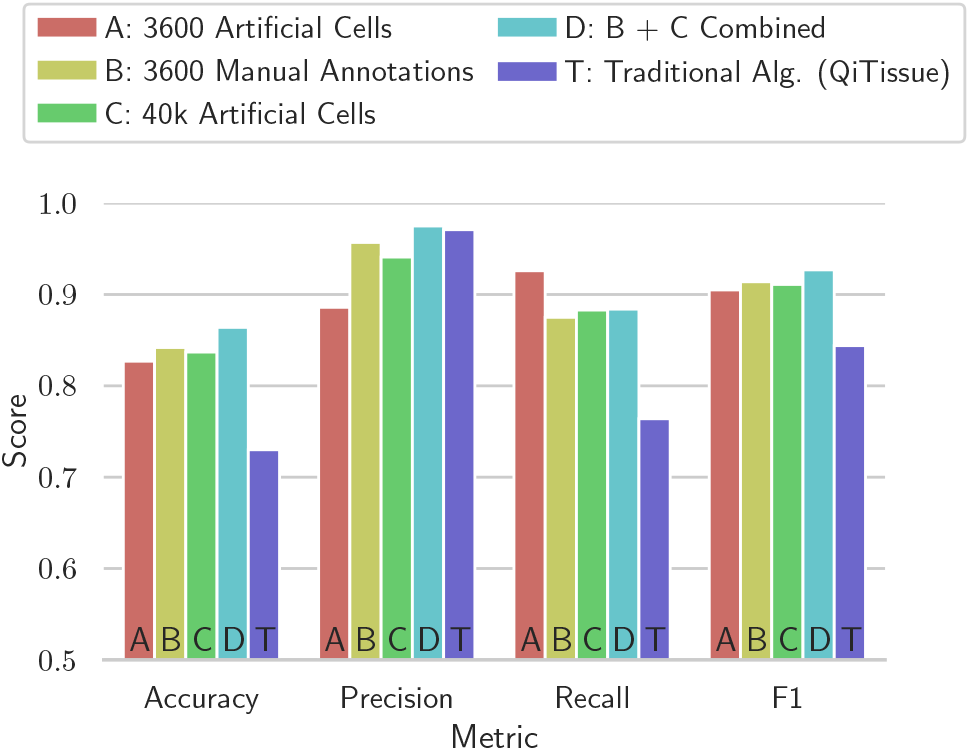
Segmentation quality of models trained with different training data and of a traditional algorithm. Evaluated on a test set containing about 3000 real cells from *Image B*.

#### L.4. Traditional Algorithm vs. Neural Network

As a representative example of an advanced, traditional algorithm, the segmentation feature of the software ‘QiTissue’ by Quantitative Imaging Systems, LLC, was used. It combines different algorithmic techniques such as adaptive thresholding, Watershed transformation and multicut.

The algorithm was compared to the best performing neural network *Model D* and tested in the same way as the other models. The parameters were set by an expert. The results in fig. 6 show that the neural network outperforms the traditional algorithm in all relevant metrics. It has a 19% higher accuracy and the F1 score is 10% better. The precision is almost the same as both algorithms show very few false positives. However, the neural network is sometimes better at not splitting large cells erroneously into two.

#### L.5. State-of-the-Art vs. Ours Trained with only Artificial Data

To see how well a model trained only with artificial data stacks up against other state-of-the-art open-source approaches, it was tested using the evaluation framework of (2). The same dataset of 200 images^2^ as in their study was segmented with our *Model C* from section L.2. The resulting segmentations were exported as labeled mask images and fed into the benchmark code of the original evaluation.

At IoU threshold 0.5, our model performed better than both the CellProfiler basic and advanced pipeline and the Deep-Cell solution, while having a lower F1 score than the U-Net model in the original comparison. Starting with IoU threshold 0.7, the F1 score of our model drops below the DeepCell score. At 0.75, the performance of our model decreases significantly and is also lower than the one of the CellProfiler and Random Forest algorithm.

This suggests that our model is universal enough to be directly transferred to other datasets, as long as the focus is on localization and not segmentation accuracy.

### M. Evaluation of Classification Results using Few And Weak Annotations

To show how this tool can be used to modify an existing instance segmentation network to also do cell phenotype classification, we classified cells into being epithelial or not. It is a good example of using manual annotations additionally, as the artificial data does not distinguish any tissue types yet.

While the ground truth data was created by leveraging CK19 and VIM stainings as weak annotations and then manually correcting the remaining errors, the challenge for the neural network will be to classify the cells only based on the nucleus shape. It will only have the DNA-marking DAPI channel as its input and no other biomarker.

*Model D* from section L.3 was used as the initial weights to train new models with a third output channel for the classification result.

Two different test sets were used to evaluate the classification performance. While both have a balanced mix of epithelial and other cells, the cells on the first one are easier to distinguish than in the second one. The results of the former are shown in fig. 7.

**Fig. 7.**
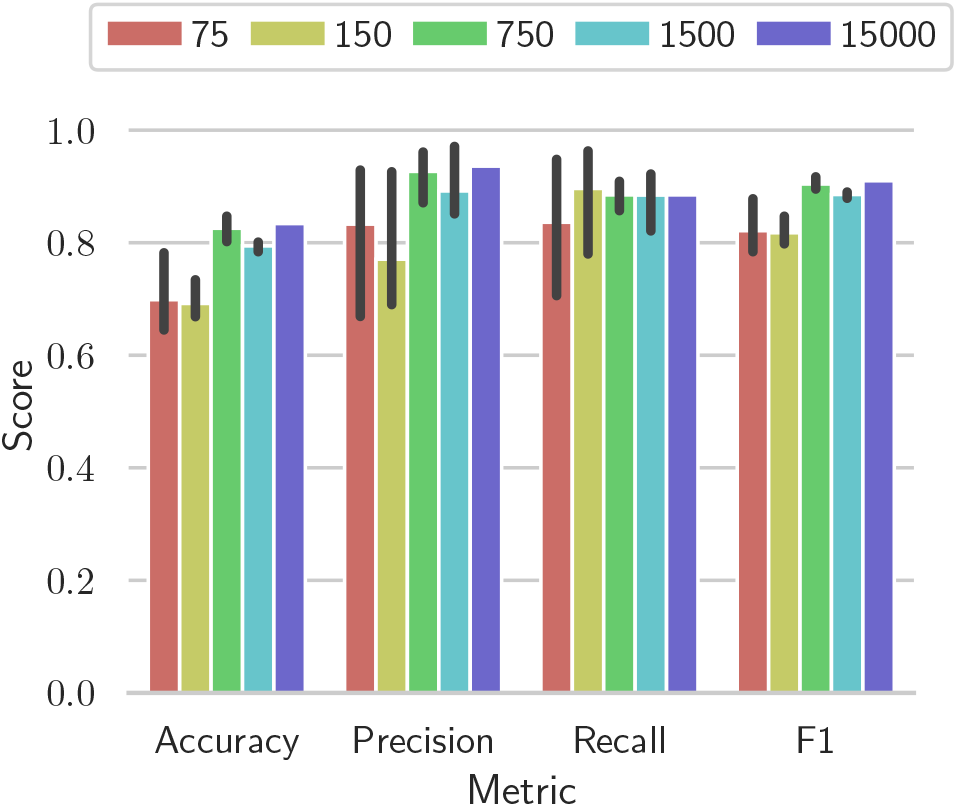
Performance of per-cell-classification into epithelial-or-not in a simple region of *Image A*. Bars show different models trained with 75 to 15000 annotated cells. From left to right for each metric: 75, 150, 750, 1500 and 15000 annotated cells in training data.

In the easier test set, even models trained with only 75 annotated cells can reach F1 scores of above 0.8. In contrast, the more difficult test set required 1500 annotations to achieve a consistent F1 score of about 0.78 across multiple runs. Still, 1500 cells can be annotated with the help of weak annotations in under an hour.

## Discussion

The evaluation of the annotation tool shows that the ‘Interactive Watershed’ assistance leads to 43% faster annotations and a semantic segmentation can speed up the process by 68%. Accurate segmentations can be achieved the fastest by first applying an instance segmentation neural network model to it and then manually correcting the result in the included annotation tool. This is six times faster than manually annotating the cells and shows how this tool can help with large scale annotations in real/world scenarios.

It was shown that artificial data is almost as good as manual annotations on real images to train a segmentation model. The more accurate boundaries of the synthetic data seem to outweigh the shortcomings of the algorithmic data generation. It suggests that the artificial data is well suited for this specific combination of cells, microscope and staining. A limitation however is, that for different input data, either the artificial data creation has to be changed, or the model has to be refined using manual annotations.

The evaluation of the amount of artificial training data shows that a model can be successfully trained with only 500 unique cells. It takes only about four minutes to prepare and generate this synthetic data from scratch and two minutes to train a network with patches cropped from it. This is faster than all the preparation times stated for other approaches in (2).

A hybrid model that was first trained using artificial data and then fine/tuned using manual annotations on real images outperformed our other models. This shows that additional manual annotations can still improve the result and a good annotation tool is beneficial for that. It also shows that transferlearning works for refining models in this way.

Compared with other state-of-the-art algorithms, our model trained without any manual annotations showed equal performance at low IoU thresholds and was slightly worse with higher IoU thresholds. As our model was not trained at all with the training data of the benchmark, this shows that the artificial data generalize well for those instance segmentation tasks where the biggest priority is the localization of the cells. The results look very similar to those reported in (25), which may hint at a characteristic of the common star-convex polygon data structure.

The lower scores of our model at higher IoU thresholds mean that the boundaries do not match well with the ground truth. Our model is therefore not suitable for tasks requiring accurate boundaries on images different from the artificial training data.

Figure 7 shows that in some cases just 75 annotated cells are enough to train a classifier. Those 75 cells can be labeled in under a minute and the training time is less than two minutes. Training a classifier to quickly identify in which regions of the image which kind of tissue dominates, is therefore possible in under three minutes. See fig. 8 for example results of such a model.

**Fig. 8.**
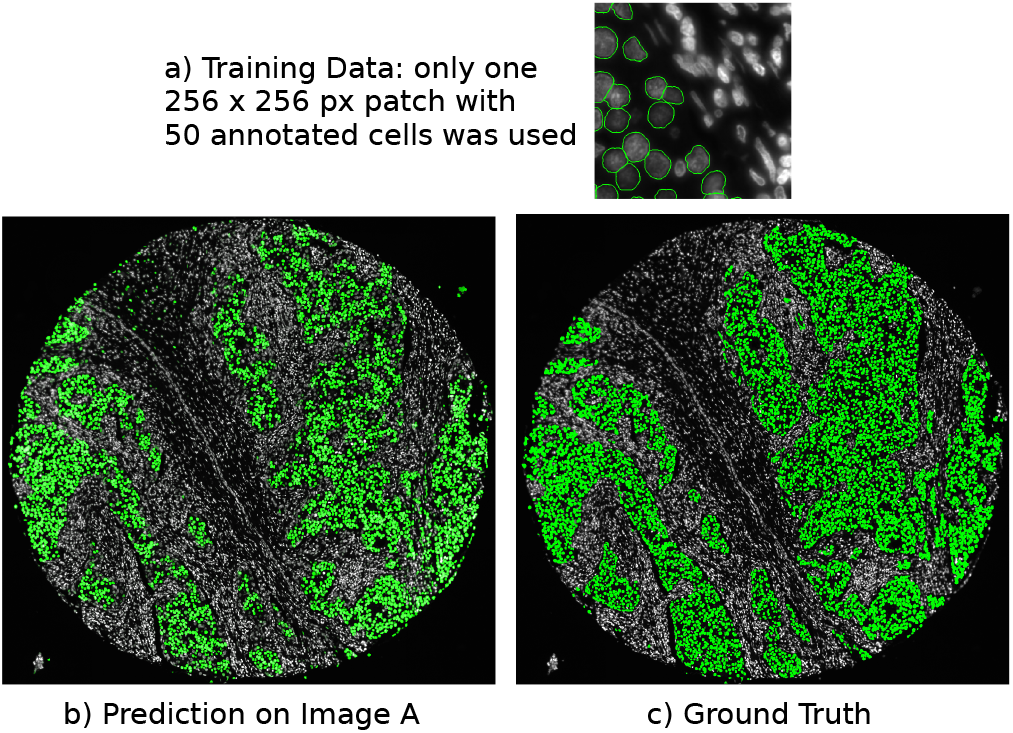
Classification results of model trained with very few annotations already show the general distribution of tissue types across the image. (a) Data used to retrain a segmentation-only model for per-cell classification. (b) Prediction of the model on *Image A*. (c) Ground truth of epithelial cells in *Image A*. (a-c) Epithelial cells are marked in green, others appear in grey.

On the other side, it is still worth to annotate a larger set of cells as in a more difficult test set only models trained with 1500 or more cells consistently give usable results.

## Conclusion

Using our tool it is possible to create working deep learning models for cell segmentation in about six minutes. Using it to annotate cells is six times faster than manually annotating them. Extending such a network with classification features can be done in three to ten minutes.

All the findings support the hypothesis that interactive deep learning for cell analysis is possible and is a valuable alternative to traditional cell analysis algorithms and shallow learning techniques.

The tool fulfills its goal to save time in annotating single large fluorescence images and opens the door to new insights by allowing to easily integrate deep learning into the analysis.

Future work could include extending the artificial data generation for different cell, tissue and imaging kinds and retrieving more parameters automatically from real example images. While our approach is more lightweight and does not need training, a GAN may be a good addition for more complicated cases.

## Acknowledgment

We like to thank the team of Quantitative Imaging Systems, LLC, for its constructive support in realizing this tool.

## Footnotes

1 https://github.com/luminosuslight/pathology-ml-model-training

2 BBBC039 from the Broad Bioimage Benchmark Collection, available at https://data.broadinstitute.org/bbbc/BBBC039/

## Notes

### Competing Interest Statement

The authors have declared no competing interest.

